# Characterization of heterologously expressed fibril filaments, a shape and motility determining cytoskeletal protein of the helical bacterium *Spiroplasma*

**DOI:** 10.1101/2021.10.30.466559

**Authors:** Shrikant Harne, Pananghat Gayathri

## Abstract

Fibril is a constitutive filament forming cytoskeletal protein of unidentified fold, exclusive to members of genus *Spiroplasma*. It is hypothesized to undergo conformational changes necessary to bring about *Spiroplasma* motility through changes in body helicity. However, in the absence of a cofactor such as nucleotide that binds to the protein and drives polymerization, the mechanism driving conformational changes in fibril remains unknown. Sodium dodecyl sulphate (SDS) solubilized the fibril filaments and facilitated fibril purification by affinity chromatography. An alternate protocol for obtaining enriched insoluble fibril filaments has been standardized using density gradient centrifugation method. Visualization of purified protein using electron microscopy demonstrated that it forms filament bundles. Probable domain boundaries of fibril protein were identified based on mass spectrometric analysis of proteolytic fragments. Presence of both α-helical and β-sheet signatures in FT-IR measurements suggests that fibril filaments consist of assembly of folded globular domains, and not a β-strand based aggregation similar to amyloid fibrils.

## Introduction

Fibril is a *Spiroplasma*-specific nucleotide-independent filament forming protein ^1,2^. It is a homo-polymer of a 59,000 Da protein called fib or fibril that is proposed to be organized as a polymeric assembly ^3^. It is a bi-domained protein, consisting of an N-terminal domain approximately 30 % similar to 5’-methylthioadenosine nucleosidase (MTAN) and a C-terminal domain of unidentified fold ^4,5^. A set of fibril polymers and MreB filaments are present on the cytoplasmic side of the cell membrane and constitutes the cytoskeletal ribbon (Figure 1A) ^6,7^. The ribbon has been proposed to confer helical shape and to function as a linear motor contributing to motility of spiroplasmas ^3,6–10^. However, the detailed molecular architecture of fibril and the cytoskeletal ribbon remains to be determined.

**Figure 1.**
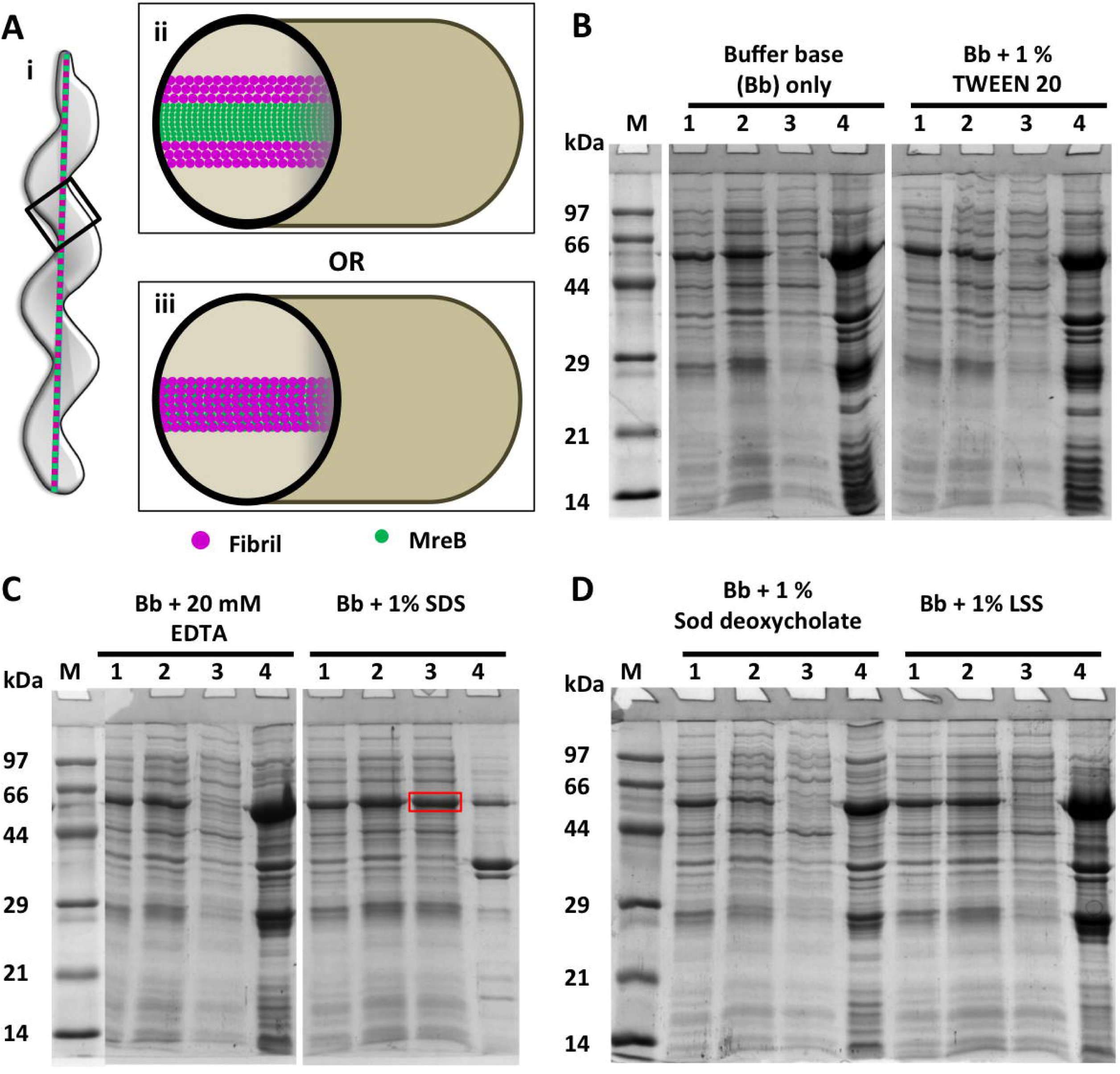
Cytoskeletal organization of *Spiroplasma* fibril and its heterologous expression and solubility check. A) Pictorial representation of intra-cellular localization of cytoskeletal ribbon of *Spiroplasma* (i) is shown. Longitudinal section (represented within the black box in (i)) of *Spiroplasma* cell showing schematic representation of fibril and MreB organization as proposed by Kürner et al., 2005 (ii) and Trachtenberg et al., 2008 (iii). According to Kürner et al., (2005), fibril filaments flank the MreB filaments while Trachtenberg et al., (2008) proposes that the MreB filaments are sandwiched between fibrils and membrane. B - D) Solubility check of fibril overexpressed in *E. coli* using different detergents [sodium dodecyl sulphate (SDS), sodium deoxycholate (Sod deoxycholate), N-lauroylsarcosine sodium salt (LSS)] and ethylenediaminetetraacetic acid (EDTA). Lanes correspond to total lysate (1), supernatant after spinning the lysate at 4,629 xg (2), supernatant (3) and pellet (4) after 159,000 xg spin of the clarified lysate. Addition of SDS (1 % w/v) helped obtain fibril soluble upon 159,000 xg centrifugation and is highlighted with a red box.

According to the current model for *Spiroplasma* motility, fibril undergoes conformational changes to bring about length variations in the fibril filaments ^3,4,8^. Co-ordinated cycles of alternate contraction and extension of the fibril filaments can cause change in handedness of the cell body that are reflected in the form of kinks which push the surrounding liquid and propel the cell forward ^3,6,9^. Indeed, low resolution electron microscopy data suggests the possibility of different conformations of fibril ^4,11^. The molecular mechanism of kink propagation based on conformational changes of a constitutive filament such as fibril is also unknown.

Apart from fibril, MreB is a constituent of the cytoskeletal ribbon of *Spiroplasma* ^6^. The role of MreB in *Spiroplasma* motility was speculated long ago ^12,13^ but was experimentally demonstrated recently ^10^. Although MreB5 (one of the 5 MreB homologs in *Spiroplasma*) binds fibril and membrane ^10^, it is currently not known if MreB links fibril to the membrane. The fact that MreB3 and MreB5 form ATP-dependent dynamic polymers *in vitro* ^14,15^ suggests their possible role as the motor which induces changes in the cytoskeletal ribbon *in vivo*. However, the organization of the cytoskeletal ribbon, consisting of MreBs and fibril filaments, inside *Spiroplasma* cell is under debate (Figure 1A) ^6,7,16^.

Recent advances in electron cryomicroscopy allow us to obtain three-dimensional reconstruction of proteins and their assemblies at atomic resolution ^17,18^. A high resolution structure of fibril may facilitate identification of conformational changes ^8^ proposed to be necessary for bringing about kinking motility of *Spiroplasma* ^5,9^. The structure may also provide insights about the evolution of fibril and the uncharacterized mode of motility driven by cytoskeletal filaments.

Here, we demonstrate that fibril expressed in *E. coli* forms filamentous assemblies which are very stable and can withstand SDS treatment. Solubilization of the filaments using SDS enabled purification of fibril polymers formed by a hexahistidine-tagged construct using affinity chromatography. Alternatively, fibril was also purified using density gradient centrifugation without the use of SDS. Visualization of fibril obtained by these protocols using electron microscope showed polymers similar to those reported from *Spiroplasma* cells ^5^. Fibril filaments and their secondary structure content remained unaffected by SDS treatment, based on observation using electron microscopy and FT-IR (Fourier Transform Infrared) spectra. Mass spectrometric characterization of proteolytic fragments confirmed a probable domain boundary for the folded fibril monomers.

The study paves the way for further characterization of fibril function using biochemical assays for studying binding of fibril with other bacterial cytoskeletal proteins such as MreBs, to probably unravel a mechanism based on filament interactions between multiple cytoskeletal proteins. Further, the heterologous expression enables us to generate suitable mutant constructs for understanding the enigma behind *Spiroplasma* motility, which was hitherto hindered because of challenges in genetic manipulation of *Spiroplasma*. The variety of approaches used for obtaining structural insights on fibril provides a workflow of options for characterization of challenging assemblies of cytoskeletal filaments.

## Material and methods

### Cloning

The *fibril* gene was amplified from genomic DNA of *S. melliferum* (DSMZ accession number 21833) using appropriate primers (Supplementary table S1). Six TGA codons (corresponding to amino acid number 26, 200, 258, 316, 323 and 342 from amino terminus) in *fibril* gene were mutated to TGG and full-length *fibril* gene with modified tryptophan codons was cloned into pHIS17 vector (Addgene plasmid #78201) with or without a terminal hexa-histidine (His_6_) tag either at the N-terminus or at the C-terminus. All the clones were confirmed by sequencing (Supplementary table S1).

### Expression

The full-length *fibril* gene with modified tryptophan codons was used for expression of fibril protein with or without a terminal (His_6_) tag in *E. coli* cells. All the fibril constructs were overexpressed in *E. coli* BL21 (AI) cells (Invitrogen) by growing the transformants in Luria Bertani (LB) broth supplemented with ampicillin (100 µg/mL final concentration) at 37 °C under shaking conditions until OD_600_ reached 0.6. The cultures were then induced with sterile L-arabinose (final concentration 0.2 %) and were grown post-induction at 25 °C for 6 hours. Protein overexpression was confirmed by the presence of high intensity band with molecular weight of ∼ 59,000 Da on the SDS-PAGE gel.

### Solubility check of fibril

During initial attempts to purify fibril the clarified lysate was spun at 159,000 xg to separate filamentous proteins from monomeric or oligomeric components. Accordingly, fibril was obtained in the pellet fraction. However, fibril from the pellet could not be solubilized and hence required screening of different compounds for their ability to solubilize fibril.

### Standardization for solubility of fibril protein

Pellet obtained from 1.2 L culture expressing His_6_-fibril was re-suspended in 270 mL of lysis buffer base [10 mM Tris (pH 7.6), 1 % (v/v) Triton-X 100 and 2 M glycerol]. Cells were lysed and the lysate was equally divided into 6 tubes. Multiple solubilization conditions for the lysate were tested with one of the following detergents added to buffer base: N-lauroyl sarcosine sodium salt (LSS; 0.5 % w/v), Sodium deoxycholate (1 % w/v), Tween-20 (1 % v/v), sodium dodecyl sulphate (SDS; 1 % w/v), EDTA (20 mM w/v). All these conditions show the effect of respective compounds in addition to Triton X-100, a constituent of lysis buffer base. The effect of only Triton X-100 (1 % v/v) was checked by addition of lysis buffer base to the sixth tube containing lysate. Values in bracket indicate final concentration of respective compounds in the 50 mL lysate. The tubes containing lysate were incubated on ice for 30 minutes and then spun at 4629 xg to remove cell debris as pellet. The supernatant was further spun at 159,000 xg at 4 °C for 30 minutes. Supernatants obtained after centrifugation at 159,000 xg were transferred to fresh tubes and each pellet was re-suspended in 2 mL T_10_E_10_ buffer [10 mM Tris (pH 7.6), 10 mM EDTA (ethylenediaminetetraacetic acid)]. 10 µL sample from each condition at all the stages was mixed with 10 µL SDS-PAGE loading dye (2X), heated at 99 °C for 10 minutes and visualized on SDS-PAGE gel. 7 µL was loaded on the gel for total lysate and 159,000 xg pellet while 15 µL was loaded for 4,629 xg supernatant and 159,000 xg supernatant.

### Purification of fibril expressed in *E. coli* by Ni-NTA affinity chromatography

A 200 mL culture pellet of *E. coli* BL21 (AI) cells expressing fibril with a terminal His_6_ tag (either N-terminal or C-terminal end) was re-suspended in 50 mL lysis buffer [50 mM Tris (pH 7.6), 200 mM NaCl and glycerol 10 % (v/v)]. Cells were lysed using a probe sonicator while incubation on ice and lysate spun at 4629 xg at 4 °C for 15 minutes to remove cell debris as pellet. Supernatant was further spun at 159,000 xg at 4 °C for 30 minutes to pellet down fibril filaments. The pellet was re-suspended using 1.8 mL buffer A_200_ [50 mM Tris (pH 7.6), 200 mM NaCl] at room temperature. 200 µL of 10 % (w/v) SDS solution was added and mixed with the re-suspended pellet. The protein suspension was spun at 21,000 xg at 25 °C for 10 minutes to remove any large insoluble aggregates. The soluble fraction from this step was spun at 159,000 xg at 25 °C for 30 minutes and fibril was obtained in the supernatant. All the steps after addition of SDS were performed at room temperature (∼ 25 °C) since SDS precipitates in cold condition (4 °C).

The supernatant was mixed with 2 mL Ni-NTA resin (Ni-NTA agarose; Qiagen) pre-equilibrated with buffer A_200_ (50 mM Tris pH 7.6, 200 mM NaCl) and left for mixing upon gentle agitation at room temperature for 30 minutes. The mixture was put into an empty column with filter. The bound protein was eluted using elution buffer containing increasing concentrations (25 mM, 50 mM, 100 mM, 250 mM and 500 mM) of imidazole in binding buffer. Fractions containing protein of interest were identified on a 12 % SDS-PAGE gel, pooled and concentrated using concentrators. The salts and imidazole were removed by performing 4 cycles of 10-fold dilution of the concentrated protein with sterile distilled water, followed by concentration. The salt-free, concentrated protein was used for visualization using Field Emission-Scanning Electron Microscope (FE-SEM) or Transmission Electron Microscope (TEM).

### Purification of insoluble fibril using density gradient

#### Preparation of urografin density gradient

Sterile urografin solution (76 % w/v; Cadila healthcare Ltd, Kundaim, India) was obtained for preparation of density gradients. To prepare a 30.4 % (w/v) solution of urografin, 125 µL Tris (2 M; pH 7.6) and 500 µL EDTA (500 mM) were added to 10 mL of urografin in a tube and volume made to 25 mL by addition of distilled water. A 53.2 % (w/v) urografin solution was prepared in a separate tube by addition of 125 µL Tris (2 M; pH 7.6) and 500 µL EDTA (500 mM) to 17.5 mL of urografin and volume made to 25 mL by addition of distilled water. Both the solutions were vortexed to ensure uniform solution in each tube. The density gradient was prepared by layering 4.5 mL of 30.4 % urografin solution on top of 4.5 mL of 53.2 % urografin solution in an ultra-clear centrifuge tube (Beckman Coulter; catalogue number 344059). The tubes were left at room temperature (25 °C) for about 20 hours to allow the formation of a linear gradient.

### Purification of fibril from *E. coli* by separation on urografin density gradient

A 500 mL culture pellet of *E. coli* BL21 (AI) cells expressing full length untagged *fibril* (UTF) gene was re-suspended in 60 mL of lysis buffer [10 mM Tris pH 7.6, 1 % (v/v) Triton X-100] and lysed using a probe sonicator. The lysate was spun at 4,629 xg at 4 °C for 10 minutes and cell debris in the form of pellet was discarded. The supernatant so obtained was further spun at 159,000 xg to separate fibril filaments (pellet) from the non-polymerized proteins (supernatant). The pellet contained enriched fibril protein and was re-suspended in 1 mL T_10_E_10_ buffer [10 mM Tris (pH 7.6), 10 mM EDTA] and left for stirring at 4 °C for about 20 hours. The stirring breaks pellet clumps into finer particles which can be separated on the density gradient.

1 mL of stirred enriched fibril was loaded on top of 9 mL urografin density gradient. The tubes containing gradient loaded with protein were spun in a swinging bucket rotor at 100,000 xg at 25 °C for 2 hours. The two protein layers were collected as separate fractions using a pipette. Urografin was removed from the fractions by two rounds of washes by diluting it with 65 mL distilled water and spinning at 159,000 xg at 25 °C for 30 minutes. The pellet obtained after second round of washing was re-suspended in sterile distilled water (500 µL) and used for visualization on 12 % SDS-PAGE gel along with marker proteins to identify fraction containing fibril.

### Visualization of fibril filaments

#### Sample preparation for visualization of fibril filaments using Field Emission-Scanning Electron Microscopy (FE-SEM)

Pieces (5 mm X 5 mm) of silicon wafer disc (Sigma-Aldrich) were thoroughly washed with 100 % ethanol and stored in 70 % ethanol at room temperature until use. For sample preparation, the silicon wafer pieces were air dried at room temperature to allow evaporation of residual ethanol and water mixture. 2-3 µL of the fibril purified using affinity chromatography was drop casted on dried silicon wafer pieces and allowed to dry in dust-free environment at 37 °C. 15 nm thick gold coating was then done on the silicon wafer pieces with completely dried protein samples, using sputter coater (Quorum Tech) before visualizing the samples under FE-SEM (ZEISS).

#### Sample preparation for visualization of fibril filaments using Transmission Electron Microscopy (TEM)

Carbon-Formvar coated copper grids (300 mesh; Ted-Pella, Inc) were used without or with glow-discharging [two rounds of 15 mA current for 25 seconds using plasma cleaner (Quorum technologies) just before use]. Samples were prepared by applying 2-5 µL of purified protein to grids and allowed to stand at room temperature for at least 30 seconds to facilitate settling down of the filaments. Excess buffer was blotted using Whatman filter paper (Sigma-Aldrich). 2-4 µL stain (0.5 % w/v uranyl acetate) solution was applied to the grid and immediately absorbed from bottom using blotting paper so as to prevent absorption of the stain by protein. The grids were allowed to dry at room temperature in a dust-free environment for at least 2 hours. Dried grids were stored in a grid storage box in a dry, dust-free environment until observation in a TEM. Grids were scanned using transmission electron microscope (JEM-2200FS, Jeol Ltd.) initially at low magnification and protein imaged at higher magnifications.

#### Fourier-transform infrared (FT-IR) spectroscopy

His_6_-tagged fibril purified using SDS and affinity chromatography followed by dialysis was flash frozen in liquid nitrogen and lyophilized under vacuum for about 48 hours. Similarly, untagged fibril purified by urografin density gradient was also lyophilized to remove traces of water. The dried protein samples were used for collecting FT-IR spectra. Sample transmittance was recorded in the IR spectra (1600-1690 cm^-1^) with a resolution of 2 cm^-1^ and 24 scans at each point were acquired in FT-IR Spectrophotometer (NICOLET 6700) using KBr pellet. The transmittance of a sample was normalized by considering its transmittance at 1600 cm^-1^ as 100 % and the transmittance at other wavelengths as the percentage of transmittance at 1600 cm^-1^. Values obtained after normalization were subtracted from 1 to obtain absorbance (as percentage) and it is plotted against wave number (cm^-1^) to identify peaks corresponding to secondary structures.

#### In-gel trypsin digestion and mass spectroscopy of the protein

Purified/enriched protein was visualized on 12 % SDS-PAGE gels by Coomassie staining. The most prominent bands (corresponding to masses ∼ 60,000 Da, ∼ 36,000 Da and ∼ 26,000 Da) were excised from the gel, cut into small pieces and transferred to fresh tubes. Gel pieces were de-stained by re-suspending in buffer containing 40 % acetonitrile and 50 mM TEABC [triethylammonium bicarbonate] for 30 minutes. Stain was removed by spinning the sample (21000 xg at 25 °C for 5 minutes) and discarding the supernatant. The de-staining procedure was repeated 5 times to ensure complete removal of Coomassie stain. Protein in gel was reduced by addition of freshly prepared 10 mM DTT (dithiothreitol) to cover gel pieces and incubation at 60 °C for 30 minutes. Excess DTT was then removed and alkylation carried out by addition of 20 mM iodoacetamide and incubation in dark for 15 minutes. Upon incubation iodoacetamide was removed and gel pieces were dehydrated by 4 cycles of washing using acetonitrile (100 %). At the end of 4^th^ cycle, acetonitrile was completely removed and completely dried gel pieces were obtained. The gel pieces were re-hydrated by re-suspension in sequencing grade trypsin (10 ng/µL in 50 mM TEABC) on ice for 60 minutes. Excess trypsin was removed without centrifugation and then gel pieces were re-suspended in 20 mM TEABC buffer overnight at 37 °C. Next, trypsin-digested peptides were obtained by spinning the samples (21000 xg at 25 °C for 5 minutes) and collection of supernatant. Remnants of the peptides were extracted from the gel pieces by re-suspending them in varying concentrations of acetonitrile: formic acid (3 % : 0.4 %, 40 % : 0.4 % and 100 % acetonitrile) followed by incubation for 10 minutes at room temperature, spinning at 10,000 xg for 10 minutes and harvesting the supernatant. The aliquots of liquid containing peptides were pooled and evaporated using CentriVap DNA vacuum concentrator until dry. The desalting and cleaning of tryptic peptides was performed using StageTip protocol ^19^.

For LC-MS/MS analysis, a Sciex TripleTOF6600 mass spectrometer interfaced with an Eksigent nano-LC 425 instrument was used. Trypsin-digested peptides (∼1 μg) were loaded onto an Eksigent C18 trap column (5 μg capacity) followed by elution on an Eksigent C18 analytical column [15 cm × 75 μm (internal diameter)] using a linear gradient of acetonitrile. A typical liquid chromatography (LC) run consisted of post-loading for 60 min onto the trap at a constant flow rate of 300 nL/minute with solvent A (water and 0.1% formic acid) and solvent B (acetonitrile). The gradient schedule for the LC run was 5 % to 10 % (v/v) B for 2 min, a linear gradient of B from 10 % to 30 % (v/v) over 50 min, 30 % to 90 % (v/v) over 4 min, 90 % (v/v) B for 5 minutes and equilibration with 5 % (v/v) B for 2 minutes. For all the samples data was acquired in information-dependent acquisition (IDA) mode over a mass range of m/z 300 − 1600. Each full MS survey scan was followed by MS/MS of the 13 most intense peptides. Dynamic exclusion was enabled for all experiments (repeat count 2; exclusion duration 5 s).

The peptides were identified out using Protein Pilot (version 2.0.1, Sciex) using Pro-Group and Paragon algorithms against the RefSeq protein database of *E. coli* (last downloaded 2021-03-01) with manual addition of fibril sequence (*S. citri*; NCBI RefSeq id WP_071937222.1) without a tag or with a C-terminal 6xHis tag separated by amino acids GS (GSHHHHHH). Iodoacetamide alkylation of cysteine as a static modification and the oxidation of methionine and N-terminal acetylation as variable modifications were defined while searching the peptides. The peptides detected from 2 different batches of each sample were pooled and non-redundant peptides marked on the fibril sequence.

## Results

### Fibril filaments withstand treatment with 1 % SDS

Having overexpressed fibril constructs in *E. coli*, we checked the fibril solubility. Fibril filaments were found in the pellet fraction upon 159,000 xg spin of the cell lysate. Re-suspension of fibril from the 159,000 xg pellet in lysis buffer did not give a clear solution, indicating that these filaments were insoluble. Thus, to obtain fibril in soluble form for its further purification, re-suspension and solubilization was attempted using different detergents (Tween 20, LSS, sodium deoxycholate, SDS, Triton X-100) and EDTA. EDTA was selected to depolymerize any nucleotide-dependent filament forming proteins such as MreB, if relevant, and prevent them from pelleting down along with fibril when spun at 159,000 xg. Only the addition of SDS at a final concentration of 1 % (w/v) was found to help in obtaining fibril in the solution (Figure 1B-D).

### Fibril purified by SDS treatment followed by Ni-NTA affinity chromatography form filament bundles

The small amount of fibril in the lysate (without addition of SDS) did not bind to the Ni-NTA matrix, probably because of occlusion of the filament bundles from the matrix or non-exposure of the hexa-histidine tag. However, the SDS-solubilized (His)_6_-fibril bound to Ni-NTA affinity matrix and could be further purified by affinity chromatography (Figure 2A). Despite the purification step, additional bands of molecular weight of about 26 kDa were consistently observed along with fibril protein band (at ∼ 59 kDa). Fibril protein thus obtained could be dialysed into water, without loss of protein through precipitation. This enabled visualization of the protein using FE-SEM. Visualization of purified fibril using FE-SEM revealed protein bundles or twisted ribbons (Figure 2B). Observation of fibril assemblies using transmission electron microscopy (TEM) revealed that these bundles (Figure 2C and D) are indeed protein filaments assemblies and appeared similar to those reported from *Spiroplasma* ^5^. It is interesting to note that fibril polymers remained unaffected despite treatment with SDS. In order to avoid use of SDS and also to ensure that filament formation was not affected by SDS treatment, an alternative protocol was standardized for purification of fibril, by use of density gradient ultracentrifugation.

**Figure 2.**
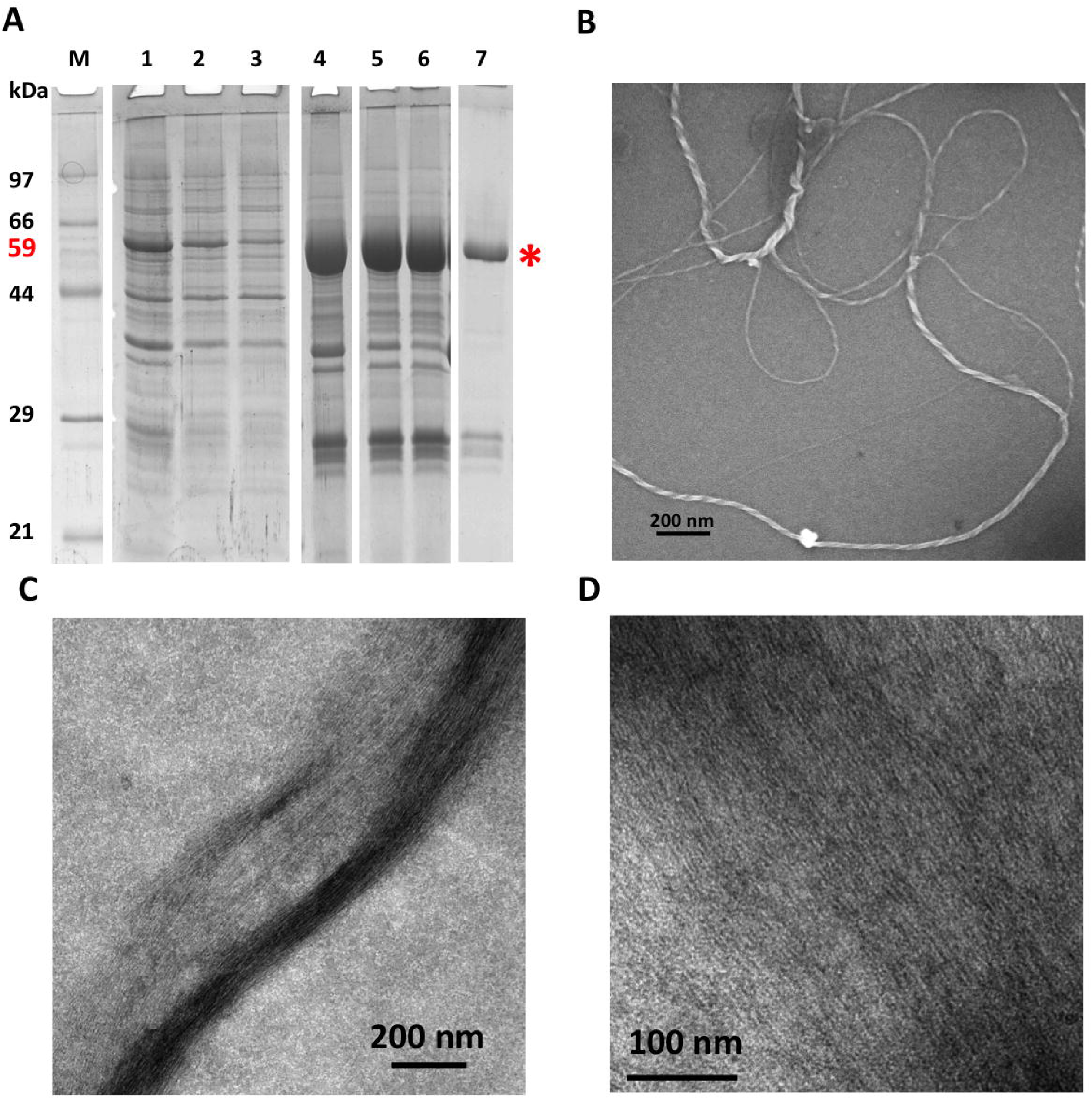
Purification of hexahistidine-tagged fibril by SDS treatment and affinity chromatography. A) A 12 % SDS-PAGE gel showing purity of fibril at different stages of purification. Lanes correspond to total lysate (1), supernatant after spinning clarified lysate at 4,629 xg (2), supernatant (3) and pellet (4) after 159,000 xg spin of the clarified lysate, supernatant of the 159,000 xg pellet solubilized using 1 % SDS (w/v) followed by spin at 21,000 xg (5), supernatant obtained after spinning the soluble protein from step 5 at 159,000 xg (6), protein fraction purified using Ni-NTA affinity chromatography (7). Scanning electron microscopy (SEM; B) and negative staining transmission electron microscopy (TEM; C and D) images of fibril filaments purified using SDS treatment followed by affinity chromatography.

### Fibril filaments purified using density gradient ultracentrifugation are present as bundles

Resuspended fibril pellet obtained after 159,000 xg ultracentrifugation separated into 2 different bands on the urografin density gradient (Figure 3A). Visualization of the protein in the two bands on SDS-PAGE gel after removal of urografin by dilution and pelleting revealed that the lower band contains enriched fibril (Figure 3A, lane 6). Electron microscopy observation of the samples revealed filament bundles similar to that obtained with SDS treatment (Figure 3B-D).

**Figure 3.**
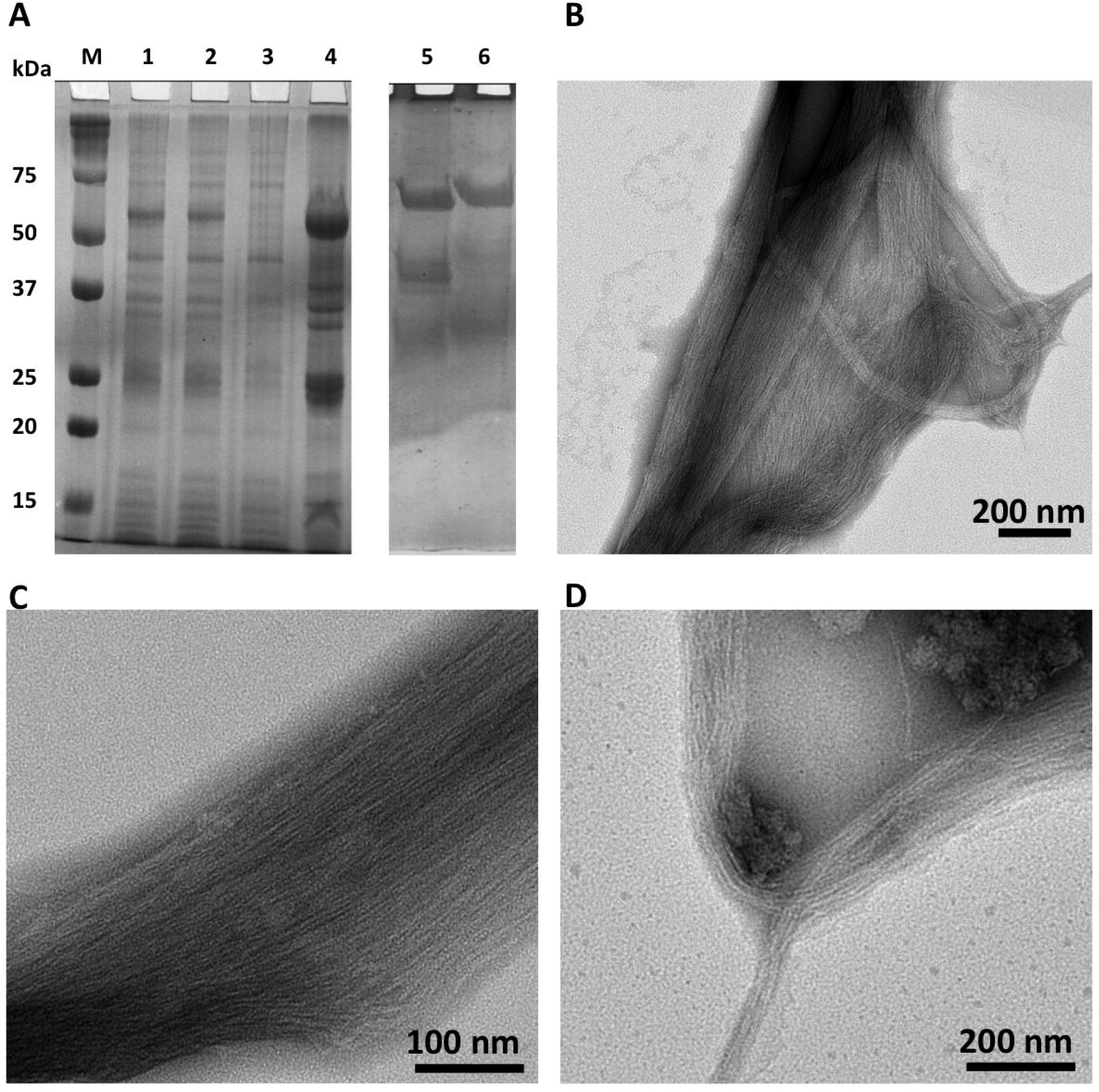
Purification of untagged fibril using urografin density gradient. A) A 12 % SDS-PAGE gel showing purity of fibril at different stages of purification. Lanes correspond to protein marker (M), total lysate (1), supernatant after spinning lysate at 8,000 xg (2), supernatant (3) and pellet (4) after spinning 8,000 xg supernatant at 159,000 xg respectively, top (5) and bottom (6) layers formed on the gradient by separation of 159,000 xg pellet fraction. Upon purification, the protein migration is retarded due to residual urografin associated with the protein. B-D) Negative staining transmission electron microscopy (TEM) images of fibril filaments purified using urografin density gradient. Individual thin filaments are visible in the images.

Next we investigated if the SDS treatment affected fibril by comparing secondary structures of fibril purified with and without the use of SDS using FT-IR spectroscopy.

### Fibril filaments exhibit secondary structure content of α-helices and β-sheets

The FT-IR spectra for fibril purified with or without SDS (using urografin density gradient) shows similar profiles (Fig 4). All the three protein constructs show the presence of peaks corresponding to α-helices (1655 cm^-1^), β-turns (1685 cm^-1^) and β-sheets (1636 cm^-1^) ^20,21^ suggesting the presence of these secondary structures in the fibril constructs purified by different protocols. Thus, based on FT-IR data we conclude that fibril is not an amyloid-like aggregate of protein, but has specific secondary structures, alpha helices and beta sheet.

**Figure 4.**
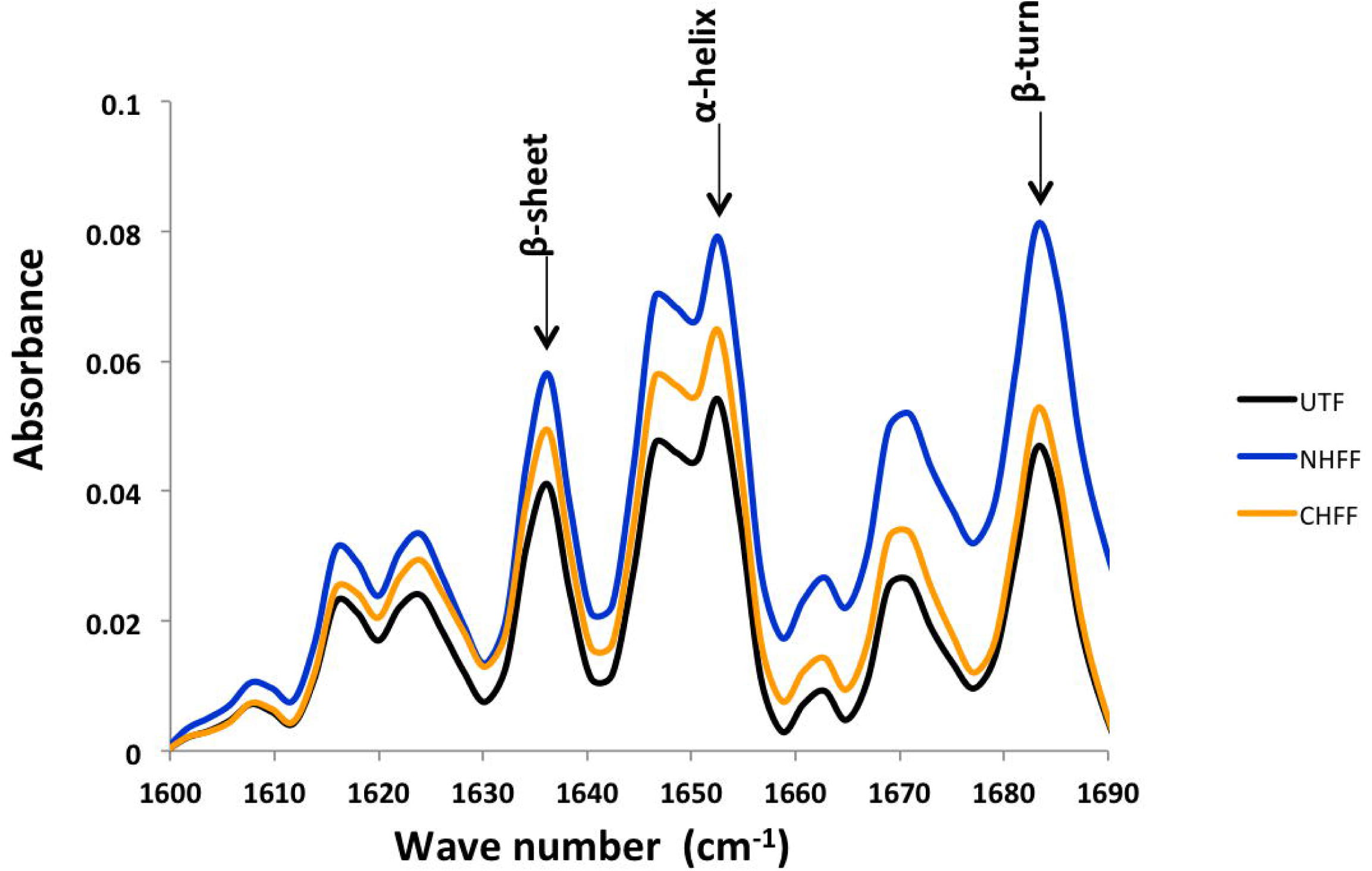
Fibril remains unaffected by SDS treatment. FT-IR analysis of untagged fibril (UTF) purified using urografin density gradient, His_6_-fibril (N-terminal His_6_ fibril; NHFF) and fibril-His_6_ (C-terminal His_6_ fibril; CHFF) purified by SDS treatment followed by affinity chromatography shows similar peak pattern. All the three protein constructs display peaks around 1655, 1636 and 1685 cm^-1^ corresponding to α-helices, β-sheets and β-turn secondary structures, respectively. This data indicates that fibril is not affected by the SDS treatment employed for its purification.

Having confirmed the stability of fibril upon SDS treatment, we decided to explore the identity of low molecular weight proteins accompanying fibril, a 59 kDa protein. The most prominent of these proteins were bands corresponding to molecular weight of ∼ 36 kDa and ∼ 26 kDa (Figure 3A, lanes 5, 6). These bands appear to be similar to those observed during purification of His_6_-tagged fibril using SDS and affinity chromatography (Figure 2A) as well as in the supernatant and pellet fractions at stages of differential centrifugation steps (Figure 2A and 3A). This led us to hypothesize that the ∼ 36 kDa and 26 kDa proteins must be either breakdown products of fibril or are contaminant proteins with the molecular weight of about ∼ 26 kDa having affinity to the Ni-NTA matrix or fibril. Thus we performed mass spectrometry study to reveal the identity of these proteins.

### Proteolytic fragments of fibril identify approximate domain boundaries of fibril

Analysis of the peptides obtained from the ∼ 36 kDa and ∼ 26 kDa proteins accompanying enriched fibril purified by density gradient and ∼ 26 kDa proteins associated with purified His-tagged fibril by in-gel trypsin digestion revealed that these are indeed breakdown products of full-length fibril (Figure 5). This suggests that fibril got proteolysed during heterologous expression in *E. coli* or the purification procedure. However, the fragments were retained with the fibril polymers most likely because they form an integral core of the fibril filament, and proteolysis occurred at the exposed loops in the polymerized state.

**Figure 5.**
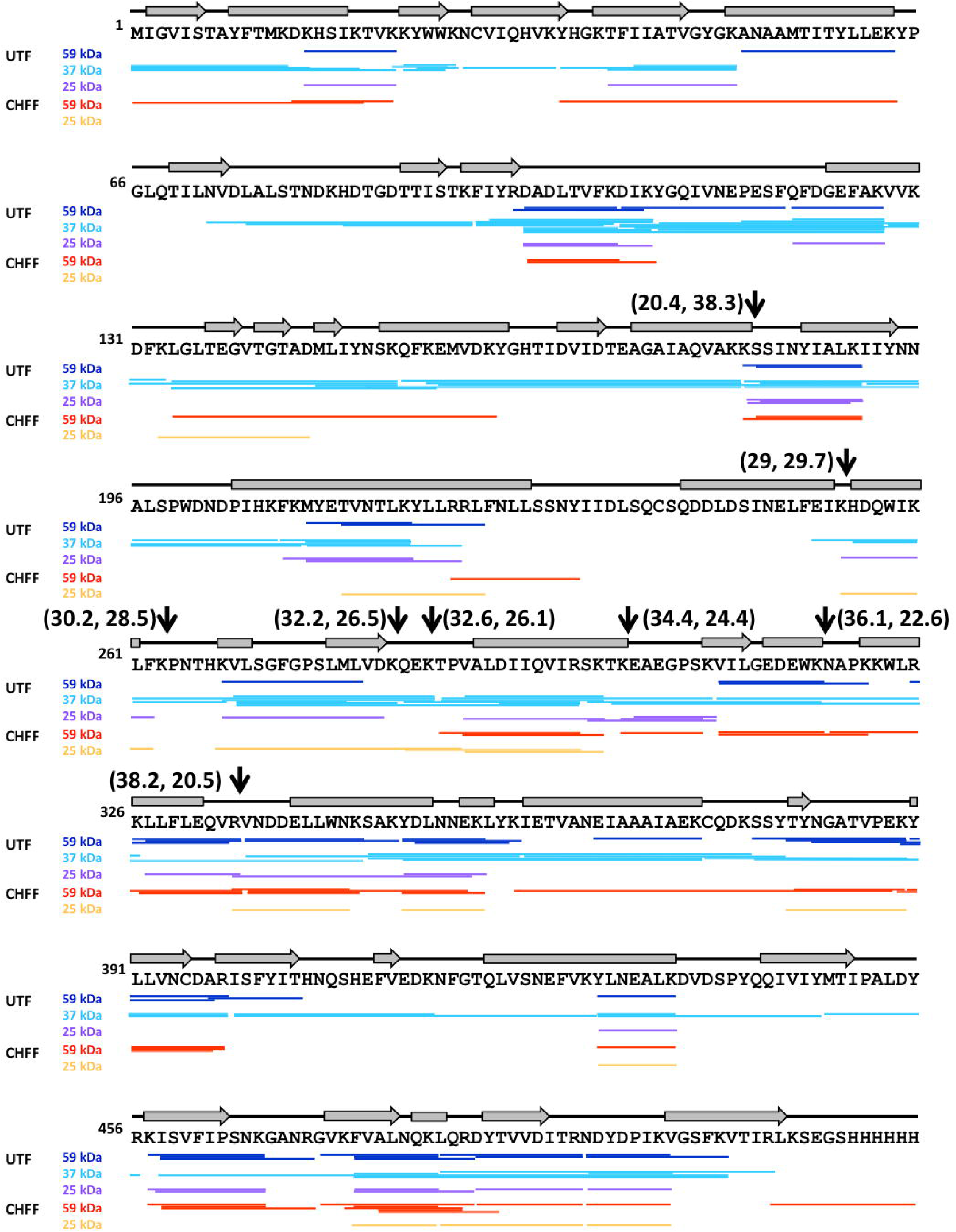
Fibril is proteolyzed in the cell. Mass spectrometry analysis of trypsin-digested peptides of about 59,000 Da, 36,000 Da and 26,000 Da bands observed during the purification of untagged fibril (UTF) and fibril with a C-terminal 6xHis tag (CHFF) by mass spectrometry study revealed that the 36,000 Da and 26,000 Da bands are indeed breakdown products of fibril. Amino acid sequences of individual peptides detected by mass spectrometer are represented by lines below the corresponding sequence while secondary structures (grey arrows-strands, grey sheets-helices) as predicted using PSIPRED (http://bioinf.cs.ucl.ac.uk/psipred/) are shown above the sequence and residue numbers are shown in superscript. Black arrows point to the potentially exposed trypsin protease target sites. Values in brackets next to black arrows indicate the masses in kilodalton (kDa) of the amino and carboxy-terminal fragment if the trypsin cleaved fibril at that position.

Analysis of fibril protein sequence revealed that its proteolysis around residues L^220^ or G^312^ can result into complementary fragments with molecular weights of ∼ 36 kDa and 26 kDa. These suggest that the peptides consisting 1-220 could constitute the amino-terminal domain while residues 312-512 make up the carboxy-terminal domain (Figure 5). The region consisting of residues 220-312 could potentially be the flexible linker connecting the two domains with each other.

## Discussion and Conclusions

The ability to control protein polymerization often proves useful for its purification ^22^. Thus actin, tubulin and their bacterial homologs were the initial cytoskeletal proteins to be purified and hence are well characterized. In contrast, intermediate filaments and other classes of constitutive filaments such as pili, flagella form stable, insoluble polymers and hence require treatment with denaturing agents like guanidine hydrochloride or urea for solubilizing them to enable their purification ^23^. Similarly, constitutive filament-forming proteins from bacteria, bactofilins and crescentin, were purified using denaturing condition (6 M urea or 6 M guanidium chloride) or using gradient centrifugation ^24–26^. Use of proteins purified by denaturation-purification-renaturation protocols for structural studies is tricky because of the risk of not knowing the correct folded state. These challenges associated with purification of constitutive filaments forming proteins have limited our understanding about their structure and function. Fibril is one such nucleotide-independent polymerizing protein classified as a cytoskeletal element ^2,27^. The role of conformational changes in fibril was hypothesized long ago ^8^ but different conformations of fibril have not yet been observed at molecular level.

During the initial attempts of purification we found that fibril remains insoluble upon pelleting at forces higher than 100,000 xg. The detergent screen revealed that fibril could not be solubilized by Tween-20, N-lauroyl sarcosine sodium salt, Triton X-100 or sodium deoxycholate but only using SDS (1 % W/V). Similar resistance of fibril has earlier been observed for 8 M urea ^28^. In addition to solubilization of fibril, the SDS treatment also exposed its 6xHis-tag thereby facilitating its purification by affinity chromatography. Since use of SDS, an anionic detergent, can denature proteins ^29^, we visualized the SDS-treated fibril to check if the protein was aggregated rather than forming filaments. To our surprise, we found that fibril was present as polymers. The observation of characteristic twists in the fibril ribbon was observed using FE-SEM. The assemblies of SDS-treated fibril are similar to those isolated from *Spiroplasma* ^5^ and suggest that fibril is resistant to SDS treatment. In order to develop a SDS-free fibril purification protocol, we used urografin density gradient to enrich insoluble fibril. Visualization of fibril purified using density gradient by TEM revealed morphologically similar assemblies to those purified using SDS as well as reported in literature ^5^.

The fibril purified by density gradient centrifugation was accompanied by low molecular weight proteins (∼ 26 kDa and ∼ 36 kDa). Also, the 6xHis tagged fibril (irrespective of the presence of tag at the N or C terminus), the ∼ 26 kDa bands remain associated with full-length fibril (59 kDa) even after purification by affinity chromatography. Mass spectroscopy analysis of these proteins accompanying fibril suggested these to be the break down products of fibril. Thus, the residues constituting ∼ 26 kDa (about 240 amino acids) from each end must constitute the two domains of fibril. Approximately 30 residues between the N and C-terminal domains must be forming a linker. This information is useful to design short constructs of fibril to identify its soluble domains, perform structural characterization and study their polymerization properties. Fibril purified with or without use of SDS showed similar FT-IR profiles re-assuring that SDS treatment did not denature fibril. The FT-IR profiles also revealed that fibril is not ‘only α-helical’ or ‘only cross-β sheets’ structure formed by α-keratin and amyloid fibrils respectively ^30,31^, but forms both α helices and well as β sheets constituting a globular protein domain.

In absence of structural information on fibril, we added the 6xHis tag at the amino or carboxy terminus of fibril to attempt its purification by affinity chromatography without affecting its polymerization. However, we observed that the fibril did not bind the Ni-NTA affinity matrix without SDS treatment. This suggested to us that the terminal 6xHis tag was not exposed and may be buried during protein folding or at the polymerization interface. The challenge of deciding the site of insertion of a tag such that the function is not affected holds true for proteins of unknown fold and is further complicated for the proteins that assemble into polymers/oligomers, as was the case of fibril. This limits the options to purify proteins of interest in high quantities, with maximum purity for biochemical and structural characterization, similar to those obtained by chromatography techniques. In such cases, density gradient centrifugation (DGC) is useful for purifying the protein of interest based on its density. The technique is widely used for the separation of macromolecular assemblies such as protein polymers, viruses and lipoproteins from other biomolecules ^32–35^. The DGC remains as the technique of choice for label-free purification of biomolecules, especially filament assemblies ^36–38^.

In summary, we have performed characterization of fibril and standardized protocols for its purification. Our approach helps obtain purified protein in quantities sufficient for structural studies and biochemical characterization of fibril using *E. coli*. Another use of the standardized heterologous expression system for fibril is its domain dissection. The lack of sophisticated tools for genetic modification of *Spiroplasma* has hindered the progress of structural characterization of full-length fibril and identification of domains by genetic screens. The genetically refractive nature of *Spiroplasma* has even prevented researchers from dissecting fibril to identify its domain boundaries and polymerization interface. In such a situation use a genetically facile organism becomes obligatory. Our demonstration of obtaining purified, folded fibril using *E. coli* paves way for expression and purification of short constructs of fibril to confirm its domain boundaries and polymerization interfaces. The system also opens up avenues for testing interaction of fibril and other *Spiroplasma* proteins such as MreB ^10^., *in vivo* using *E. coli* by co-localization studies using fluorescent tags.

## Acknowledgements

This work is supported by funds from Department of Science and technology (DST) INSPIRE Faculty Fellowship (IFA12/LSBM-52), Innovative Young Biotechnologist Award (BT/07/IYBA/2013), Department of Biotechnology Membrane Structural Biology Programme Grant (BT/PR28833/BRB/10/1705/2018) and funds from IISER Pune to P. Gayathri. SH acknowledges IISER Pune and Infosys foundation (IISER-P/InfyFnd/Trv/1) for the fellowship. Electron microscopy facility at IISER Pune (supported by DST-Nanomission (EMR/2016/003553), proteomics facility funded by Department of Science and Technology Fund for Improvement of S&T Infrastructure (DST-FIST; SR/FST/LSII-043/2016) to IISER Pune Biology department and associated staff at IISER Pune is acknowledged. Preliminary electron microscopy experiments were carried out at the electron microscopy facility at the Indian Institute of Science Bangalore. Mr. Anil Prathamshetty, Physics department, IISER Pune, is acknowledged for techincal help with FE-SEM experiments. Authors also acknowledge Prof. Jayant Udgaonkar for the access to mass spectroscopy facility, Dr. Harish Kumar and Mr. Kundan Kumar of IISER Pune for their technical help for mass spectrometry sample preparation and analysis.

---

**Supplementary table 1.**
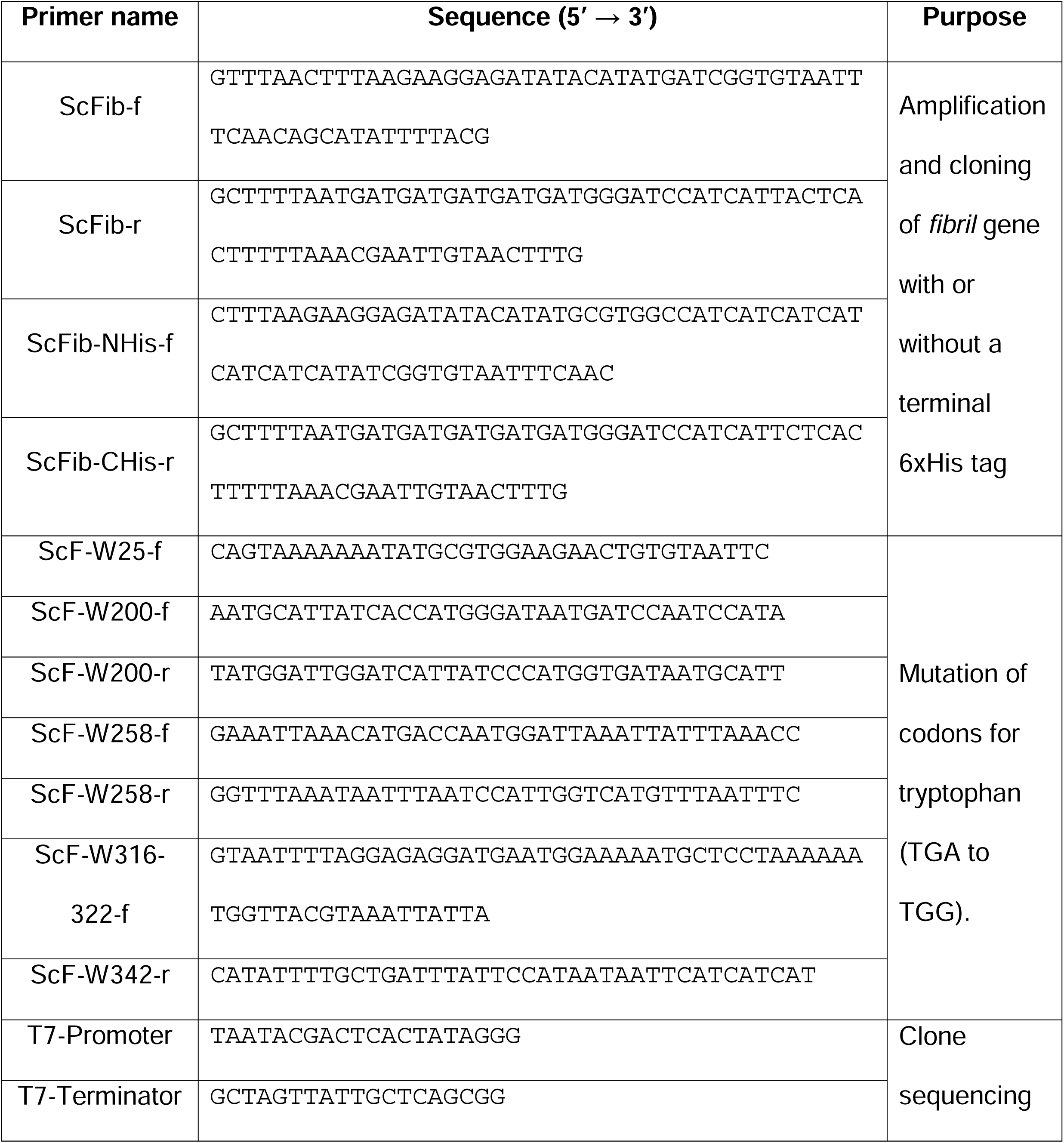
List of primers used for amplification and cloning of *fibril* gene into pHIS17 vector.

